# A Heart-on-a-Chip Microdevice with Aligned Fibers for Cardiotoxicity Assessment

**DOI:** 10.64898/2026.04.30.721826

**Authors:** Kozue Murata, Mosha Abulaiti, Rio Okama, Kensuke Kato, Yo Tanaka, Hidetoshi Masumoto

## Abstract

**Background and Objectives:** Cardiovascular cells differentiated from human induced pluripotent stem cells (iPSCs), including cardiomyocytes, are valuable for evaluating human cardiac pharmacology and toxicity. Early assessment of cardiotoxicity, especially for novel drugs like anticancer agents, is essential for improving drug development efficiency and reducing costs. This study aimed to develop a highly sensitive bioassay system capable of evaluating the physiological function of human cardiac tissue in vitro.

**Methods:** Human iPSCs were differentiated into cardiovascular cell types (cardiomyocytes, vascular endothelial cells, and vascular mural cells) and assembled into a cardiac tissue model on aligned fiber device. This tissue was cultured dynamically to induce the formation of vascular network-like structure. By combining the fiber device with our previously developed heart-on-a-chip microdevice (HMD), we created a new model of HMD (Aligned Fiber-based HMD; AF-HMD) with improved throughput and stability. Pulsatile force changes induced by drug exposure were quantified by tracking the displacement of fluorescent microbeads within the microchannels.

**Results:** AF-HMD demonstrated functional responses to known cardiac agonists and toxicants, such as doxorubicin. The device also replicated clinically relevant cardiotoxic events, including the synergistic effects of trastuzumab and doxorubicin, showing marked reductions in contractile force and beat rate, mirroring clinical observations.

**Conclusions:** The AF-HMD system provides a sensitive and reproducible platform for evaluating cardiotoxicity in drug development. It offers a promising tool for preclinical screening, with potential applications in personalized medicine and predicting cardiotoxic risk in cancer therapy.

## Introduction

Among the challenges in modern pharmacotherapy, drug-induced cardiotoxicity continues to hinder the development and clinical use of both novel and established therapeutics(1). Several drugs have historically been withdrawn from the market due to unforeseen cardiac adverse effects, despite passing conventional preclinical safety assessments such as hERG channel assays and animal-based models(2). These events have revealed critical shortcomings in traditional screening systems and highlight the urgent need for predictive, human-relevant platforms to evaluate cardiac safety(3,4). Human induced pluripotent stem cells (iPSCs) have emerged as a powerful tool for disease modeling and drug screening(5). Differentiation of iPSCs into cardiomyocytes and other cardiovascular lineages has enabled the construction of three-dimensional (3D) cardiac tissue models. Compared with monolayer cultures and animal models, these engineered tissues better recapitulate the multicellular architecture and contractile dynamics of human myocardium, providing a more physiologically relevant platform for pharmacological testing(6). To realize the potential of iPSC-derived cardiac tissues in practical drug evaluation, we developed a functional bioassay platform capable of sensitively quantifying contractile performance. The heart-on-a-chip microdevice (HMD), an organ-on-a-chip system designed to mimic cardiac function under controlled microenvironmental conditions, enables real-time quantification of parameters such as pressure, contractile force, and stroke volume(7). This is achieved by tracking the displacement of fluorescent microbeads within microfluidic channels, driven by the contraction of the engineered cardiac tissues. Nevertheless, the first-generation HMD faced critical limitations in operational stability and throughput. Specifically, tissue detachment from the push bars during long-term culture and limited sampling capacity impeded broader applicability in high-throughput or prolonged testing scenarios. To overcome these limitations, we have utilized an aligned fiber-based cell culture scaffold to develop a structurally enhanced version of the device, termed AF-HMD. This upgraded system offers improved mechanical robustness and enables cardiotoxicity assessment following relatively long-term drug exposure. In the present study, we describe the development and validation of the AF-HMD as a novel in vitro platform for cardiotoxicity screening. By integrating a dynamic rocking culture system that promotes tissue maturation with a redesigned microfluidic architecture, AF-HMD enables reproducible, quantitative assessment of pharmacological responses in engineered cardiac tissues. Furthermore, we demonstrate that this system can recapitulate clinically relevant cardiotoxic responses, including the synergistic effects observed during co-treatment with trastuzumab and doxorubicin.

## Methods

### Maintenance of Human iPS cells (iPSCs) and Differentiation of Cardiovascular cell lines

The present study used Human iPSCs lines 201B6 established at the Center for iPS Cell Research and Application (CiRA), Kyoto University, Kyoto, Japan(5). The maintenance of human iPSCs and differentiation of cardiovascular cells was conducted in accordance with our previous studies with modifications(7–10) In brief, iPSCs were expanded and maintained with StemFit AK02N medium (AJINOMOTO, Tokyo, Japan). At confluence, the cells were dissociated with TrypLE Select (Thermo Fisher Scientific, Waltham, MA, USA), dissolved in 0.5 mM ethylenediaminetetraacetic acid in PBS (1:1) and passaged as single cells (5,000 – 8,000 cells/cm^2^) every 7 days in AK02N containing iMatrix-511 silk (FUJIFILM Wako Pure Chemical Corp., Osaka, Japan) (0.125 µg/cm^2^) (uncoated laminin fragment5) and ROCK inhibitor (Y-27632, 10 µM, FUJIFILM Wako). For cardiovascular cell differentiation, single iPSCs were seeded onto Matrigel-coated plates (1:60 dilution) at a density of 300,000–400,000 cells/cm^2^ in AK02N with Y-27632 (10 µM). At confluence, the cells were covered with Matrigel (Corning, Corning, NY, USA) (1:60 dilution in AK02N) one day before induction. We replaced the AK02N medium with RPMI + B27 medium (RPMI 1640, Thermo Fisher; 2 mM L-glutamine, Thermo Fisher; 1× B27 supplement without insulin, Thermo Fisher) supplemented with 100 ng/mL Activin A (R&D, Minneapolis, MN, USA) (differentiation day 0; d0) and 5 µM CHIR99021 (Tocris Bioscience, Bristol, UK) was added for 24h, which was followed with supplementation with 10 ng/mL bone morphogenetic protein 4 (BMP4; R&D) and 10 ng/mL basic fibroblast growth factor (bFGF; FUJIFILM Wako) (d1) for 4 days without culture medium change. At d5, the culture medium was replaced with RPMI1640 medium supplemented with 50 ng/ml of vascular endothelial cell growth factor (VEGF)165 (FUJIFILM Wako), 2.5 µM IWP4 (REPROCELL, Beltsville, MD, USA) and 5 µM XAV939 (Merck, Kenilworth, NJ, USA). The culture medium was refreshed with RPMI1640 supplemented with 50 ng/ml VEGF every other day. Beating cells appeared at d11 to d15. In this protocol, we could exclusively induce cardiomyocytes (CMs) and vascular endothelial cells (ECs). To control the percentages of vascular mural cells (MCs) sufficient to form cell sheets, we used a part of differentiation culture for MC differentiation and induced the differentiation of MCs as required; after d3, the culture medium was replaced with RPMI + FBS medium [RPMI1640, 2 mM of L-glutamine, 10% fetal bovine serum (FBS)] and was refreshed every other day.

### Flow cytometry

Flow cytometry was conducted in accordance with our previous study with modifications(11,12). Differentiated cardiovascular cells and cardiac tissue sheets were dissociated by incubation with Accutase (Nacalai Tesque, Kyoto, Japan) and stained with one or a combination of the following surface markers: anti-PDGFRβ conjugated with phycoerythrin (PE), clone 28d4, 1:100 (BD, Franklin Lakes, NJ, USA) for MCs, and anti-VE-cadherin conjugated with phycoerythrin (PE), clone 55-7h1, 1:100 (BD) for ECs. To eliminate dead cells, cells were stained with the LIVE/DEAD fixable Aqua dead cell staining kit (Thermo Fisher). For cell surface markers, staining was carried out in PBS with 5% FBS. For intracellular proteins, staining was carried out in cells fixed with 4% paraformaldehyde (PFA) in PBS. Cells were stained with the anti-cardiac isoform of troponin T (cTnT) (clone 13-11) (Thermo Fisher) labelled with APC using Zenon technology (Thermo Fisher) (1:50) for CMs. The staining was performed in PBS with 5% FBS and 0.75% saponin (Nacalai Tesque). The stained cells were analyzed by CytoFLEX S (Beckman Coulter, Brea, CA, USA). Data was collected from at least 10,000 events. Data was analyzed with CytExpert software (Beckman Coulter). The percentage of CMs, ECs and MCs in cardiac tissue sheets used in the present study was as follows: CM, 52.55±10.11 %, EC, 16.23±5.71 %, MC, 18.15±5.62% (n=26) (Supplementary Figure 1).

**Figure 1:**
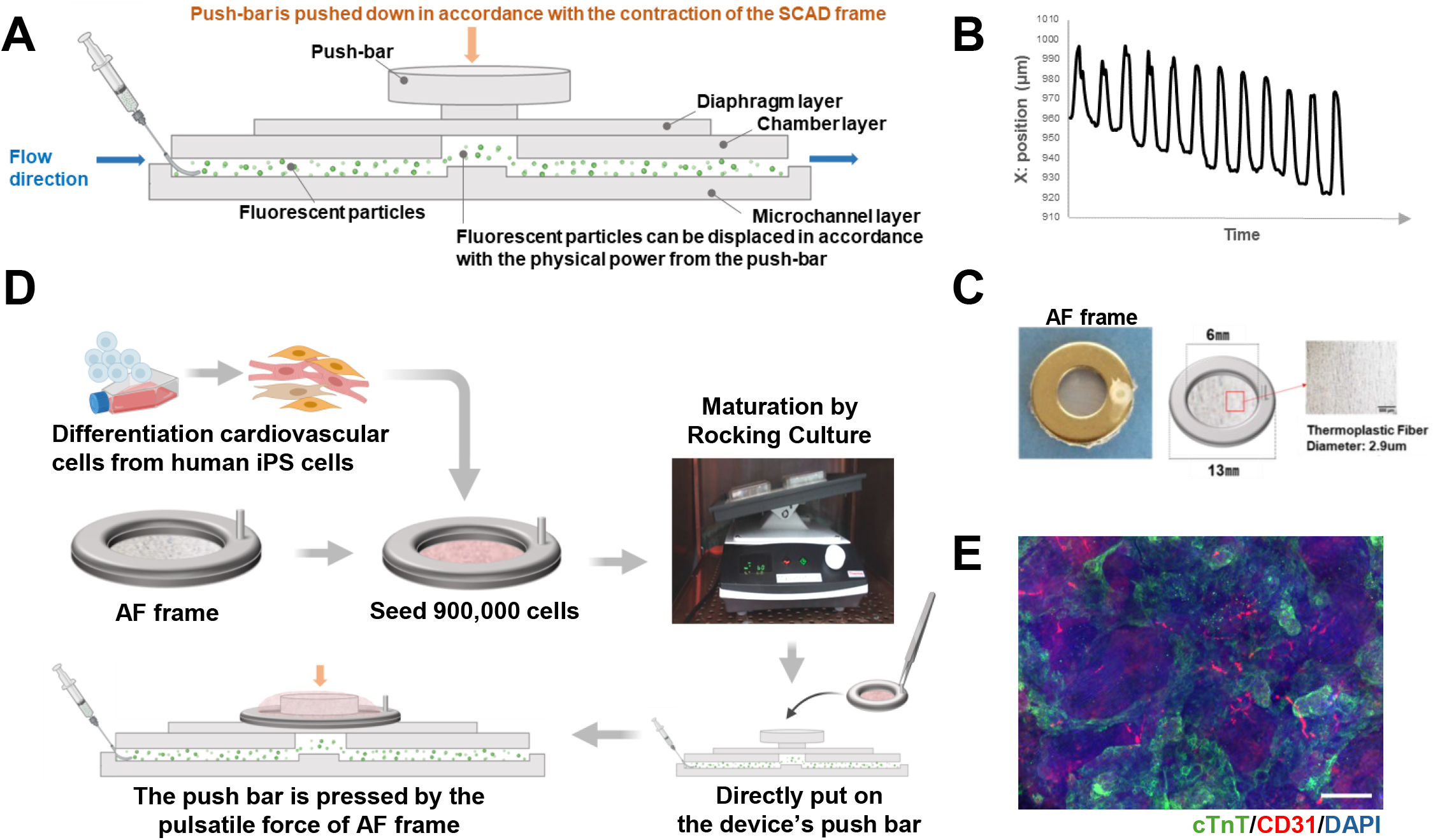
Design and Structural Overview of the AF-HMD. (A) Schematic representation of the basic structure of the Heart-on-a-chip Micro Device (HMD). (B) Representative displacement image of micro fluorescent particles embedded within the microchannel, visualizing tissue contraction-induced movement. (C) Photograph of the AF frame with a central thermoplastic fiber serving as a tissue scaffold. (D) Workflow for the fabrication and functional measurement using the AF-HMD system (E) Fluorescence immunostaining of cardiac microtissue cultured on the AF frame. Endothelial cells were stained with anti-CD31 (red), Cardiomyocytes were stained with an anti-cardiac troponin T (cTnT) antibody (green), and nuclei are counterstained with 4’,6-diamidino-2-phenylindole(DAPI) (blue). *Scale bar = 200 μm*.

### Fabrication of the AF-HMD Device

HMD (Heart-on-a-MicroDevice) was fabricated using polydimethylsiloxane (PDMS) through standard soft lithography as previously reported(7). This manufacturing was carried out by Fukoku Bussan Co., Ltd (Tokyo, Japan). To briefly describe the structure, the device consisted of a multilayered microfluidic chip containing a deformable diaphragm, a microchannel layer, and an integrated push bar. Components were assembled using oxygen plasma bonding. The aligned fiber-based cell culture scaffold frame (AF frame) was fabricated by the Stem Cell & Device Laboratory Inc. To briefly describe the structure, a solution of a thermoplastic polyether ester elastomer (Hytrel® 3001; Toray-DuPont, Tokyo, Japan) dissolved in an organic solvent was loaded into a syringe mounted on a fiber sheet fabrication system. The solution was continuously delivered for a predetermined duration onto a rotating drum collector wrapped with a substrate sheet, resulting in the formation of a fibrous sheet with an average fiber diameter of 2.9 µm. The fibers exhibited preferential alignment along the direction of drum rotation. A ring-shaped metal frame (outer diameter: 13 mm; inner diameter: 6 mm) with adhesive applied to its reverse surface was affixed to the fibrous sheet. The metal frame together with the adherent, directionally aligned fibers was then excised using a hole punch to produce an in vitro scaffold construct. To facilitate handling and maintain orientation during subsequent experiments, a silicone tube approximately 5 mm in length was attached to the metal frame along the axis of fiber alignment and served as a handling grip. Differentiated cardiac-related cells (CMs, ECs, MCs) were mixed in appropriate ratios and seeded at 900,000 cells/frame onto the AF frames. with 2 mL of attachment medium [AM; alpha minimum essential medium (αMEM) (Thermo Fisher) supplemented with 10% FBS and 5 × 10^−5^ M of 2-mercaptoethanol] containing 10 µM of Y-27632. The seeded AF frames were cultured in AM on a Compact Digital Rocker (DLAB, Beijing, China) at 60 rpm and 13 degrees for 11-14 days. We named the improved HMD system using this AF frame as the AF-HMD.

### Functional Assessment via particle Displacement Analysis

The measurement of beating force generated by cardiac tissue on AF frame (pulsatile force) was performed by modifying a previously reported method(7): the AF Frame was stabilized by its own weight on the push bar of the HMD. Using a fluorescence microscope (CKX53, OLYMPUS, Tokyo, Japan), we acquired videos of particle movement using a fluorescent isothiocyanate (FITC) filter and camera system (DP27, OLYMPUS) and software (cellSens, OLYMPUS). To facilitate accurate particle tracking, we used Move-tr/2D, a graphical user interface (GUI) software developed by Library Co., Ltd. (Tokyo, Japan). This software enabled the removal of most background noise from the images, allowing for accurate particle tracking. The x-y coordinates of the particle positions at each time point were recorded in an Excel file, and the changes in the x-y coordinates were detected as the pulsatile force generated by the heart tissue on the AF Frame using coordinate analysis software developed in-house (CellNet Co., Ltd., Tokyo, Japan). Motion amplitude was used as the primary parameter for analysis. The pulsatile index per minute (PIPM) quantifies the change in pulsatile force per minute and is calculated by integrating beats per minute (BPM) with pixel displacement(8). Conceptually, PIPM was established as an in vitro surrogate of cardiac output (CO) in clinical cardiology, where CO is defined as heart rate multiplied by stroke volume (CO = BPM × stroke volume). By incorporating both contraction amplitude and beating frequency, PIPM reflects functional output more comprehensively than BPM alone and therefore represents a physiologically relevant parameter that more closely approximates clinical cardiac performance.

### Pharmacological Testing Protocol

Pharmacological response tests were conducted using the compounds shown in **Table 1**. Heart rate and pulsatile force were measured as drug exposure increased over time or concentration. AF frames were immersed in AM containing the set concentration of drug and processed on a Digital Rocker at 60 rpm and 13 degrees at 37°C (5% CO_2_ concentration). The processing time was fixed at 20 minutes when measuring concentration-dependent responses. In each AF frame, pulsatile force was detected without drug treatment and used as a baseline. Changes in PIPM and BPM due to drug response were detected after exposure and evaluated as relative values compared to baseline values.

**Table 1:** Details of Chemicals Used.

| **Purpose** | **Product Name (Trade Name)** | **Active Ingredient** | **Manufacturer / Source** | **Catalog No. / NHI Code** | **Dose ($\mu$ M)** |
| --- | --- | --- | --- | --- | --- |
| **Negative control** | Aspirin | Acetylsalicylic Acid | Sigma-Aldrich (Merck) | A2093 | 0.1/1/10 |
| **Common cardiovascular drug** | Nifedipine | Nifedipine | RIKEN Natural Products Depository | N/A | $10^{-3}/10^{-2}/10^{-1}/1$ |
| **Known cardiotoxic compound** | Fintepla® | Fenfluramine Hydrochloride | RIKEN Natural Products Depository | N/A | 10/30/90 |
| **Known cardiotoxic compound** | Adriamycin® | Doxorubicin | Sigma-Aldrich (Merck) | D1515 | 1/10/31.6/100 |
| **Anti-cancer agent** | HERCEPTIN® for Intravenous Infusion | Trastuzumab | Chugai Pharmaceutical Co., Ltd. | 4291406D3021 (NHI Code, Japan) | 10 |

### Immunofluorescence analysis (IFA)

For IFA, cardiac tissue formed on AF frames were stained with cTnT antibody (Thermo Fisher) (1:250), CD31 (monoclonal mouse IgG1, clone 9G11) (R&D) (1:250), with DAPI (4’,6-diamidino-2-phenylindole) (Thermo Fisher) (1:1000). Anti-mouse Alexa 546 (Thermo Fisher), anti-rabbit Alexa 488 (Thermo Fisher) were used as secondary antibodies. The tissues were photographed with an all-in-one fluorescence microscopic system, BZ-X800E (Keyence, Osaka, Japan) Combined Z-stack and sectioning functions.

### Statistical Analysis

All quantitative data are presented as mean ± standard deviation (SD), unless otherwise indicated. Statistical analyses were performed using GraphPad Prism (version 9.0, GraphPad Software, San Diego, CA). Group differences were analyzed using the Kruskal–Wallis test, followed by Dunn’ s post hoc test with Holm adjustment for multiple comparison. All experiments were performed with a minimum of three independent biological replicates (n≧3), and individual sample sizes are indicated in figure legends.

## Ethical Approval

This study did not involve any personally identifiable samples, and therefore, ethical review by the institution was deemed unnecessary.

## Results

### Structural and Functional Validation of the AF-HMD System

The AF-HMD is an enhanced platform built upon the fundamental microfluidic architecture of the previously reported HMD system(7), incorporating structural optimizations to improve operational stability and the precision of functional assessment (Figure 1A). In brief, the HMD consists of a push bar, a diaphragm layer, a chamber layer, and a microchannel layer. Fluorescent particles within the microfluidic channels exhibit synchronized movement along the X-Y axis in response to the vertical displacement of the push bar. The X-Y displacement of these particles is captured as a waveform, with the magnitude of displacement represented by the amplitude in the image diagram (Figure 1B). As previously reported, both the amplitude and pulsation interval of this waveform correlate with the pulsatile force generated by actual cardiac tissue(7). The waveform peaks correspond to the end-diastolic and end-systolic phases, respectively. To improve operational stability and experimental throughput, we developed the AF frame, which features a central thermoplastic fiber that serves as a scaffold for tissue attachment. As shown in Figure 1C, this design enables more consistent placement onto the push bar compared to conventional setups, resulting in robust functional measurements that are less dependent on operator skill. An overview of the measurement scheme integrating the HMD system with the AF frame (AF-HMD) is shown in Figure 1D. After seeding cardiovascular cells onto the AF frame and subjecting them to dynamic culture, spontaneous and synchronized contractions across the entire frame began to appear approximately 4days later. After 11-14 days of maturation treatment, immunofluorescence staining for the endothelial marker CD31 confirmed the formation of vascular network-like structures within the cardiac tissue cultured on the AF frame (Figure 1E). In the AF-HMD, samples can be returned to the culture environment except during measurements for cardiotoxicity assessment. As a result, in a continuous three-day culture experiment, the control group maintained a stable PIPM throughout the observation period. Furthermore, no tissue collapses or structural instability on the push bar, which had been observed with the conventional HMD, was detected during the measurement period (Supplementary Video 1).

To assess the capability of the AF-HMD system in detecting pharmacological responses of cardiac tissue, acetylsalicylic acid (commercially known as aspirin) was employed as a negative control(13,14), while nifedipine, a calcium channel blocker, was used as a representative cardiotropic agent. The analytical performance of the system was further examined using a quality control index (QC Index), defined as the cumulative change in PIPM across aspirin concentrations, with the PIPM value immediately prior to aspirin administration normalized to 1.0. AF frames with a QC Index below a prespecified threshold were considered to satisfy quality control criteria and were included in subsequent analyses (Supplementary Figure 2A). When all AF frames were analyzed without QC Index–based filtering, aspirin produced no significant change in PIPM across concentrations (Kruskal-Wallis test, p = 0.832), confirming its suitability as a negative control. Under the same unfiltered conditions, nifedipine exhibited a decreasing trend in PIPM; however, this effect did not reach statistical significance (Kruskal-Wallis test, p = 0.074) (Supplementary Figure 2B). In contrast, when analyses were restricted to AF frames meeting the predefined quality criterion (QC Index ≦1.5), aspirin again showed no significant concentration-dependent effect (Kruskal–Wallis test, p = 0.655), whereas nifedipine induced a statistically significant concentration-dependent reduction in PIPM (Kruskal-Wallis test, p = 0.031) (Figure 2A). These findings demonstrate that QC Index–based frame selection enhances the sensitivity and analytical robustness of the AF-HMD system. By excluding low-quality frames, the system more reliably detects pharmacologically relevant, concentration-dependent responses, thereby strengthening its applicability to cardiovascular research and potential clinical evaluation.

**Figure 2:**
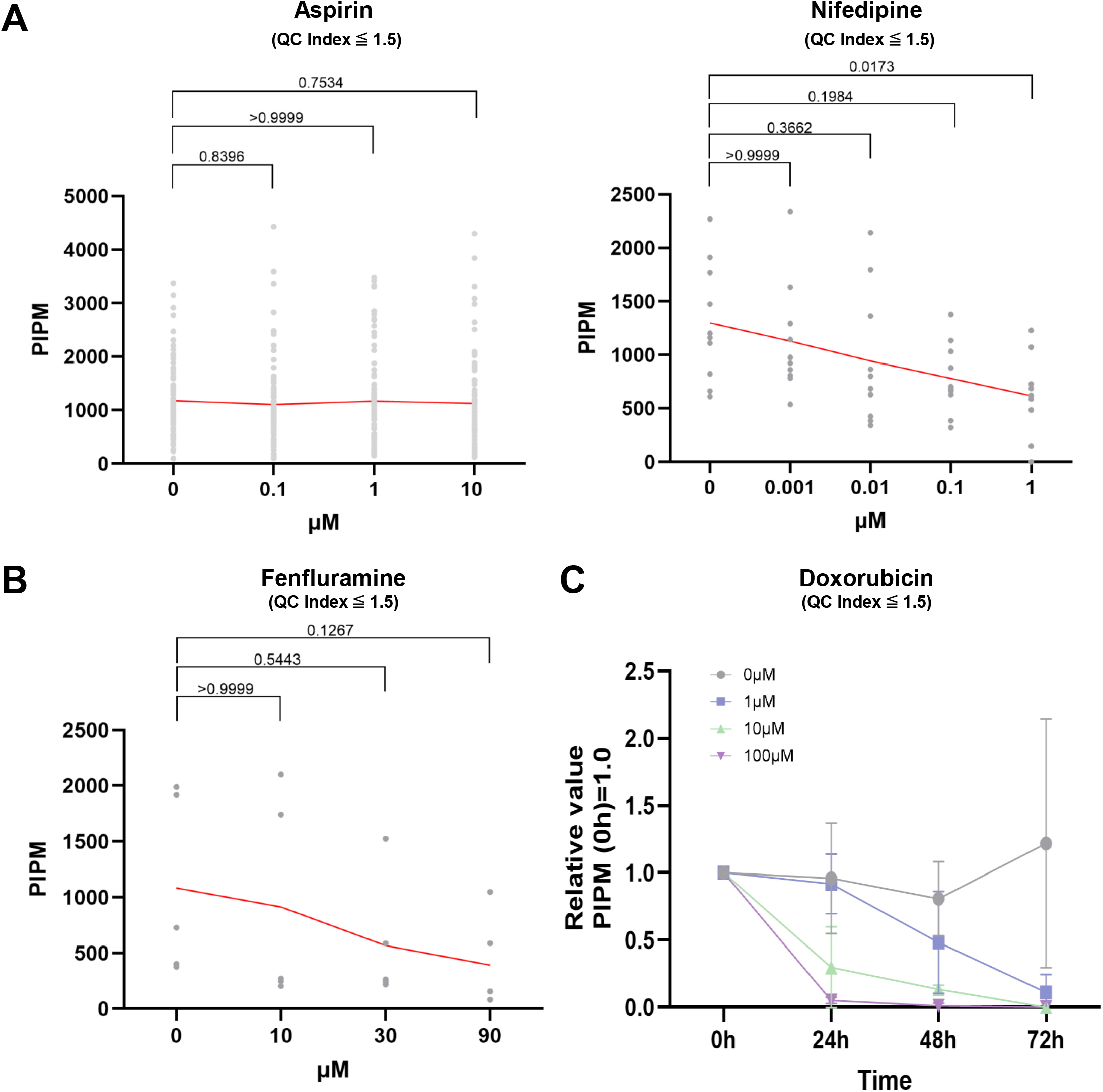
Detection of Pharmacodynamic Responses to Cardiotoxic Drug by AF-HMD. (A) Pulsatile force of micro cardiac tissues on the AF frame following exposure to the negative control compound Aspirin at concentrations ranging from 0.1 to 10 μM (QC Index≦1.5, *n=*79, Left panel). Red lines indicate the average response. Dose-dependent effects of the cardiotonic agent Nifedipine (0.001–1 μM) on Pulsatile force of AF frame-cultured cardiac microtissues (QC Index≦1.5, *n=*10, Right panel). Red lines indicate the average response. (B) Short-term (20-minute) responses to the known cardiotoxic agent Fenfluramine at concentrations of 10–90 μM (QC Index≦1.5, *n=*5). Red lines indicate the average response. (C) Time-dependent effects of the cardiotoxic compound doxorubicin (1–100 μM) on tissue contractile force following 24–72 hours of dynamic culture under shaking conditions (QC Index≦1.5, *n*=3-5). Red lines represent the average values at each concentration and time point. Cardiac output was derived from particle displacement distance multiplied by BPM, indicating the pulsatile force per minute of micro tissue on AF frame.

### Pharmacological Response to Cardiotoxic Agents

The ability to detect cardiotoxicity in humans during the early stages of drug development is critical for reducing both development time and associated costs. To evaluate the utility of the AF-HMD system in cardiotoxicity assessment, we examined its responsiveness to two well-characterized cardiotoxic agents: fenfluramine (trade name: Fentepra®) and doxorubicin (trade name: Adriamycin®). Exposure to increasing concentrations of fenfluramine resulted in a gradual reduction in PIPM (Figure 2B). In contrast, short-term exposure to doxorubicin did not elicit a concentration-dependent decrease in contractility (Supplementary Figure 2). However, during extended culture for up to 72 hours, doxorubicin treatment induced a time-dependent reduction in PIPM, which became apparent 24 hours post-treatment. Moreover, the rate of PIPM decline was accelerated in a dose-dependent manner (Figure 2C) (Supplementary Videos 1,2). These findings demonstrate that the AF-HMD system exhibits both sensitivity and specificity in detecting drug-induced cardiotoxic effects. Furthermore, the ability of the AF frame to support long-term culture underscores its potential for identifying compounds that exert delayed cardiotoxicity during prolonged exposure.

### Recapitulation of Clinically Observed Cardiotoxic Synergy

In recent years, in addition to the anthracycline-based anticancer agent doxorubicin, the monoclonal antibody trastuzumab (commercial name: HERCEPTIN® for Intravenous Infusion) has been widely employed in the treatment of HER2-positive breast cancer. It is well established that concomitant administration of trastuzumab and doxorubicin can result in cardiotoxicity. To mitigate this risk, clinical protocols have been devised that separate the timing of administration and utilize multiple treatment cycles(15,16).

To assess whether the AF-HMD system can reproduce this clinically relevant cardiotoxic interaction, cardiac tissues were treated with 10 μM doxorubicin and 1 μM trastuzumab, either as monotherapies or in combination, and the time-dependent changes in PIPM were monitored. Notably, co-treatment with both agents induced a pronounced reduction in beating rate and PIPM as early as 24-hour post-treatment. At 48 hours post treatment, cardiotoxicity in the 10 μM doxorubicin group, which was not detectable by BPM alone, became evident when assessed by PIPM. In contrast, treatment with trastuzumab alone had no significant effect on PIPM, even after 72 hours, with values remaining comparable to those of untreated control (Figure 3). The observed synergistic impairment exceeded the effects of doxorubicin monotherapy and closely recapitulates the clinical scenario in which trastuzumab exacerbates doxorubicin-induced myocardial injury. These findings demonstrate that the AF-HMD system can model complex, clinically relevant cardiotoxic phenotypes, thereby highlighting its utility for mechanistic investigations and translation cardiotoxicity assessment.

**Figure 3:**
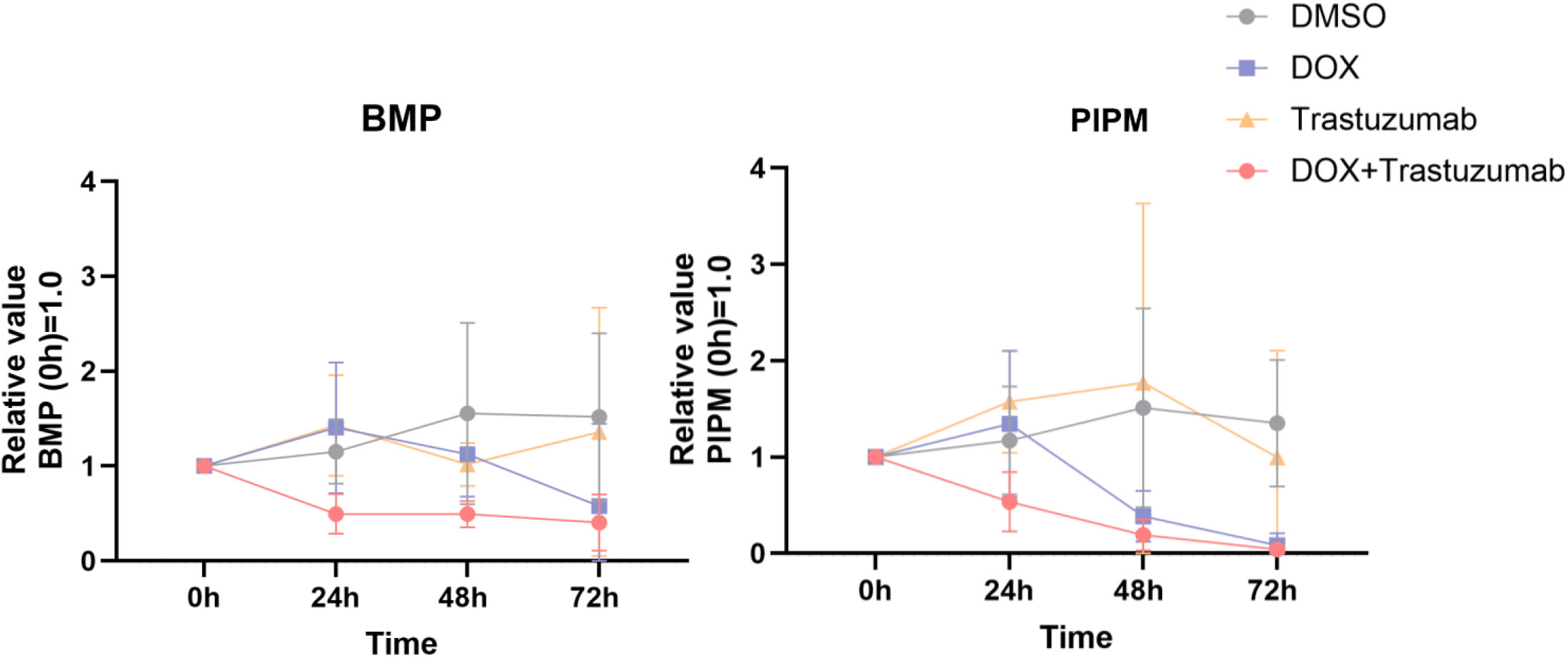
Reproduction of Clinically Observed Synergistic with Trastuzumab and Doxorubicin. (Left panel) Beating rate (beats per minute, BPM) and (Right panel) cardiac output of human iPSC-derived cardiac microtissues cultured on the AF frame following exposure to trastuzumab (10 μM), doxorubicin (1 μM), or their combination. Cardiac output was calculated as the product of particle displacement and BPM, representing pulsatile force per minute. Data are presented as mean ± standard deviation (QC Index≦1.5, *n*=3-6).

## Discussion

In this study, we established the AF-HMD platform as a physiologically relevant, robust, and reproducible in vitro system for evaluating cardiotoxicity using three-dimensional cardiac microtissues derived from human iPS cells. By preserving tissue adhesion and sustained contractile activity under long-term dynamic culture, AF-HMD overcomes a key limitation of earlier heart-on-a-chip microdevice technologies. The platform enables real-time, high-resolution monitoring of functional parameters such as contractile force and stroke volume, which are directly linked to clinically meaningful indicators of cardiac performance.

A particularly noteworthy aspect of AF-HMD is its ability to recapitulate the clinically validated synergistic cardiotoxicity induced by combined trastuzumab and doxorubicin therapy under in vitro conditions. Such adverse cardiac events have conventionally been identified only during or after clinical administration, thereby exposing patients to a substantial risk of cardiac dysfunction. In contrast, the AF-HMD platform enables the detection of these complex cardiotoxic liabilities at the preclinical stage, underscoring its potential to fundamentally transform current paradigms of cardiac safety assessment.

More importantly, the use of AF-HMD suggests a paradigm shift from establishing anticancer treatment guidelines based on retrospective clinical observations of adverse events to proactively designing therapeutic regimens and dosing strategies that minimize cardiotoxicity using an organ-on-a-chip–based alternative evaluation system. This approach allows for the anticipation and mitigation of cardiac risk without exposing patients to harm and holds particular significance in the field of cardio-oncology, where balancing oncologic efficacy with cardiovascular safety remains a critical unmet need.

In addition to recapitulating key aspects of human myocardial physiology and drug responses, the AF-HMD platform is inherently extensible. Future incorporation of patient-specific iPSC-derived cardiac tissues could enable the prediction of individual susceptibility to cardiotoxicity, thereby supporting the development of personalized therapeutic strategies. Such adaptability underscores the potential of AF-HMD not only as a preclinical screening tool but also as a translational platform for individualized risk stratification in cancer therapy. Collectively, these features position AF-HMD as a foundational technology with substantial relevance for advancing precision cardio-oncology.

## Conclusion

In this study, we developed and validated AF-HMD, an advanced heart-on-a-chip platform integrating vascularized human iPSC-derived cardiac microtissues within a microfluidic system that enables dynamic culture and functional assessment. The device faithfully reproduces essential aspects of human cardiac physiology and pharmacological responses, including clinically relevant cardiotoxic interactions. AF-HMD therefore represents a robust and versatile tool for translational cardiac safety evaluation, with broad applicability in oncology drug development and beyond.

## Limitations

Despite its strengths, the AF-HMD platform has several limitations. First, the use of iPSC-derived cardiac tissues, while advantageous for human relevance and scalability, entails a degree of cellular immaturity compared to adult myocardium, potentially limiting its predictive power for chronic or late-onset cardiotoxicity. Second, the platform currently emphasizes mechanical parameters such as contractility and beat rate. While essential, these measures may be insufficient for capturing the full spectrum of cardiotoxic effects. Integration with electrophysiological and metabolic sensors would enable a more holistic assessment of cardiac function. Third, while the system successfully modeled the synergistic effects of trastuzumab and doxorubicin, broader validation across additional drug classes and patient-specific cell sources is necessary to confirm generalizability. Lastly, AF-HMD is currently limited to a cardiac-only environment, without accounting for systemic processes such as hepatic metabolism or hormonal modulation. Integration into multi-organ-on-a-chip systems may help address this and improve systemic translational fidelity.

## Perspective

Cardiotoxicity continues to pose a significant challenge in cancer therapy, especially in the context of combination regimen. Existing preclinical models often fail to predict human cardiac responses with sufficient accuracy. AF-HMD fills this gap by offering a human iPSC-based, structurally and functionally optimized platform that can detect both individual and synergistic drug-induced cardiotoxic effects. Its capacity for real-time monitoring of contractile dynamics renders it particularly valuable for modeling complex pharmacodynamic interactions. Looking ahead, the platform’ s adaptability for use with patient-derived cells opens the door to precision medicine applications. Further developments, including integration into multi-organ systems and chronic exposure models, will enhance its predictive power. As regulatory frameworks increasingly acknowledge the value of organ-on-chip technologies, AF-HMD is well positioned to contribute meaningfully to next-generation drug safety evaluation and individualized therapeutic planning.

## Abbreviations

CVD: Cardiovascular disease
iPSC: Induced pluripotent stem cell
HMD: Heart-on-a-chip microdevice
CM: Cardiomyocyte
EC: Endothelial cell
MC: Mural cell

## Acknowledgments

We thank the members of RIKEN BDR Clinical Translational Program (Masumoto Lab), the Department of Cardiovascular Surgery at Kyoto University Hospital, Stem Cell & Device Laboratory Inc., FUKOKU BUSSAN CO.,LTD., and RIKEN Program for Drug Discovery and Medical Technology Platforms, for their collaboration and technical support.

## Figure Legends

**Supplementary Figure 1:**

**Population of cardiac structural cells on AF frame**

The proportion of each cardiac cell type within each AF frame was measured by flow cytometry. Each gray point corresponds to one frame.

**Supplementary Figure 2:**

**Quality Control Method for AF Frames**

(A) Schematic diagram of the QC Index calculation method. The QC Index was defined as the sum of the changes in PIPM at each concentration, with PIPM before aspirin administration set to 1.0. Frames were deemed to meet quality control standards when the QC Index was 1.5 or lower. (B) Pulsatile force of micro cardiac tissues on the AF frame following exposure to the negative control compound Aspirin at concentrations ranging from 0.1 to 10 μM (all frames, *n=*116, Left panel). Red lines indicate the average response. Dose-dependent effects of the cardiotonic agent Nifedipine (0.001–1 μM) on Pulsatile force of AF frame-cultured cardiac microtissues (all frames, *n=*15, Right panel). Red lines indicate the average response.

**Supplementary Figure 3:**

**Detection of Pharmacodynamic Response to Short-term Doxorubicin Treatment by AF-HMD**

Short-term (20-minute) responses to the known cardiotoxic agent Doxorubicin at concentrations of 1–100 μM (QC Index≦1.5, *n=*11). Red lines indicate the average response. Cardiac output was derived from particle displacement distance multiplied by BPM, indicating the pulsatile force per minute of micro tissue on AF frame.

**Supplementary Video 1: Representative movie of beating Cardiac tissue on AF Frame during a 3-day consecutive cardiotoxicity assessment (Control: DMSO-treated group) Supplementary Video 2: Representative movie of beating Cardiac tissue on AF Frame during a 3-day consecutive cardiotoxicity assessment (10 *μ*M Doxorubicin-treated group)**

## Central Illustration

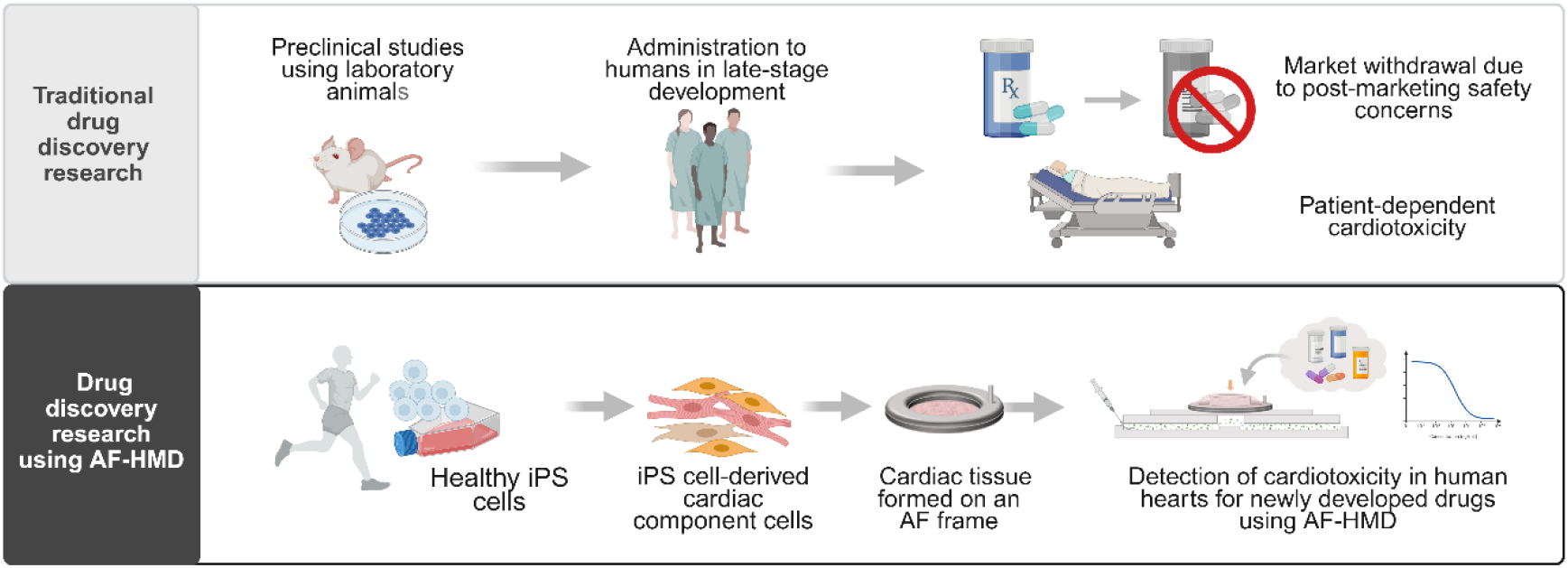

## Notes

**Funding:** This work was supported by research funding provided by a Grant-in-Aid for Scientific Research (C) (#21K08861) from the Japan Society for the Promotion of Science (JSPS), Japan (to K.M.), Stem Cell & Device Laboratory Inc. (to H.M.), RIKEN Gap Fund (to H.M.), and RIKEN BDR Organoid Project (to Y.T. and H.M.).

### Competing Interest Statement

The authors have declared no competing interest.

