## Supplementary figures and images for "A Heart-on-a-Chip Microdevice with Aligned Fibers for Cardiotoxicity Assessment"

### FigureS1

#### Supplementary Figure 1

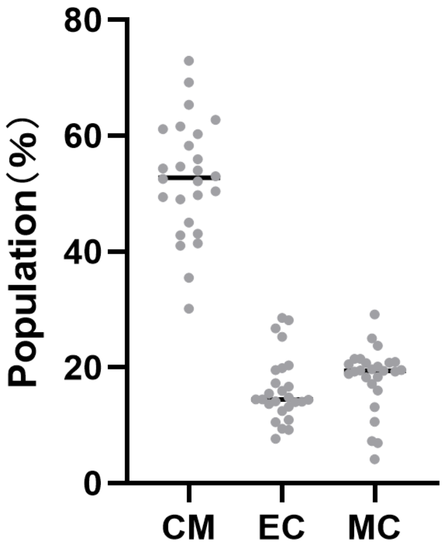

### FigureS2

#### Supplementary Figure 2

A

QC Index : Sum of change values (a+b+c)

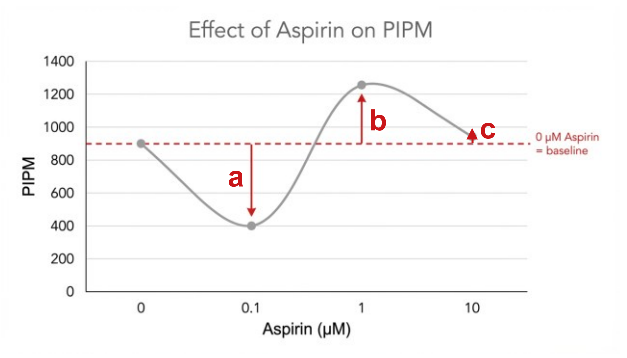


B

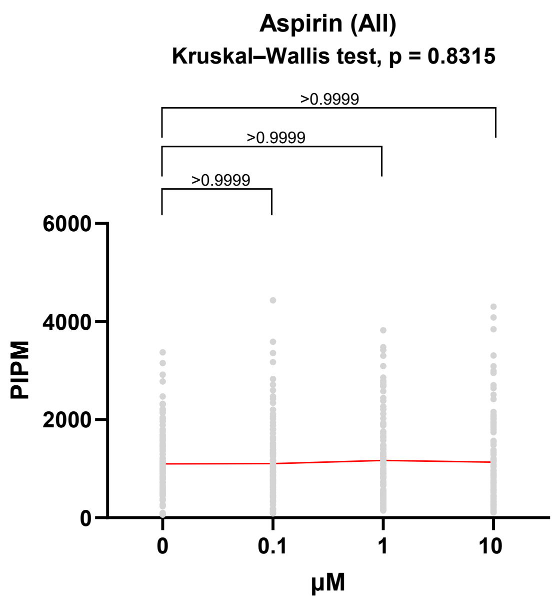


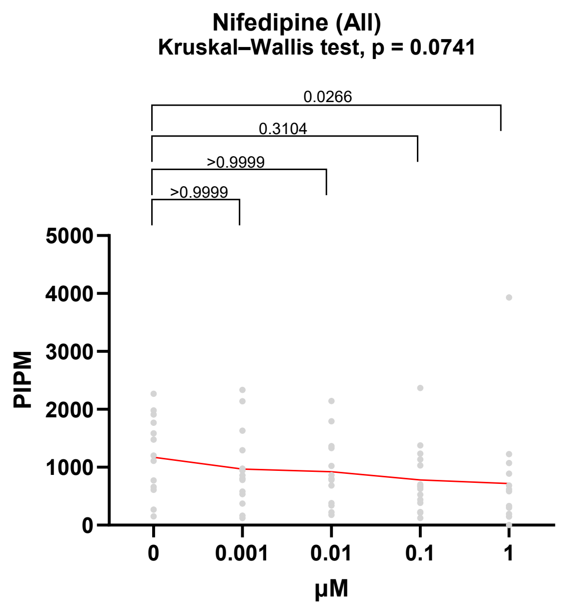

### FigureS3

#### Supplementary Figure 3

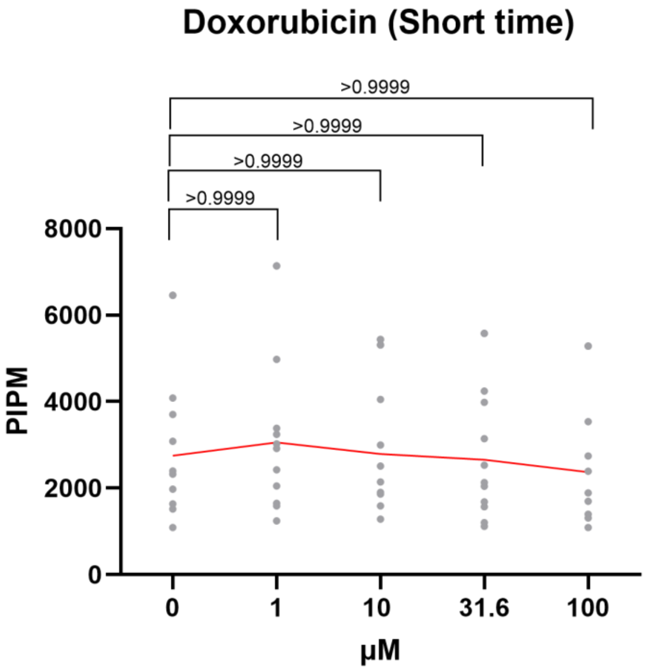
